# The movement of a leaf-derived mobile *AGL24* mRNA specifies floral organ identity in Arabidopsis

**DOI:** 10.1101/2020.11.16.385153

**Authors:** Nien-Chen Huang, Huan-Chi Tien, Tien-Shin Yu

**Affiliations:** Institute of Plant and Microbial Biology, Academia Sinica, Taipei, 11529, Taiwan

**Author notes:** **Corresponding author:** Tien-Shin Yu, Institute of Plant and Microbial Biology, Academia Sinica, Taipei, 11529, Taiwan. **One Sentence Summary**: The leaf-expressed *AGL24* mRNA moves long-distance to floral meristem to serve as a template for protein translation. The author responsible for distribution of materials integral to the findings presented in this article in accordance with the policy described in the Instruction for Authors is: Tien-Shin Yu.

## Abstract

Cell-to-cell and inter-organ communication play pivotal roles in synchronizing and coordinating plant development. In addition to serving as templates for protein translation within cells, many mRNAs can move and exert their function non-cell-autonomously. However, because the proteins encoded by some mobile mRNAs are also mobile, whether the systemic function of mobile mRNAs is attributed to proteins transported distally or translated locally remains controversial. Here, we show that Arabidopsis *AGAMOUS-LIKE 24* (*AGL24*) mRNA acts as a leaf-derived signal to specify meristem identity. *AGL24* is expressed in both apex and leaves. Upon floral meristem (FM) transition, apex-expressed *AGL24* is transcriptionally inhibited by APETALA1 (AP1) to ensure FM differentiation. The leaf-expressed *AGL24* can act as a mobile signal to bypass AP1 inhibition and revert FM differentiation. Although *AGL24* mRNA is expressed in leaves, AGL24 protein is rapidly degraded in leaves. In contrast, *AGL24* mRNA can move long distance from leaf to apex where the translocated *AGL24* mRNAs can be used as templates to translate into proteins. Thus, the movement of *AGL24* mRNA can provide the developmental plasticity to fit with environmental dynamics.

## Introduction

Cell-fate determination is shaped by the interplay between intrinsic and extrinsic signals. In plants, the shoot apical meristem (SAM) is usually surrounded by leaf primordia, which insulates SAM from direct exposure to the environmental stimuli. To ensure that development can synchronize with environmental dynamics, plants recruit various mobile molecules for inter-organ communication. During meristem transition, SAM is transformed from the vegetative meristem (VM) to inflorescent meristem (IM). Subsequently, the flanks of IM are converted into the floral meristem (FM), which differentiates to all floral organs. The switch of VM to IM and FM is reversible and highly sensitive to environmental changes. When floral-committed plants are transferred to non-floral inductive conditions, FM may generate abnormal flowers or turn back to IM or VM growth, a phenomenon known as floral reversion (Buckhout, 1880; Battey and Lyndon, 1990; Tooke et al., 2005). In many plants, such as *Impatiens balsamina*, floral reversion is affected by systemic signals derived from leaves (Pouteau et al., 1997); however, the molecular identity of leaf-derived systemic signals is largely unknown. In Arabidopsis, the meristem transition of VM to IM is controlled by a pair of antagonized mobile signals, florigen and antiflorigen, which are encoded by *FLOWERING LOCUS T* (*FT*) and *ARABIDOPSIS THALIANA CENTRORADIALIS* (*ATC*) belonging to the same gene family (Corbesier et al., 2007; Huang et al., 2012). Although it has been shown that *FT* may be involved in preventing floral reversion (Müller-Xing et al., 2014; Liu et al., 2014), it appears that the leaf-derived signals in FM differentiation may not be simply due to the movement of *FT*, because floral reversion is not usually observed in *ft* or other florigen defective mutants. Thus, the molecular identity of leaf-derived signals in controlling floral organ development remains to be elucidated.

In Arabidopsis, floral organ development is divided into multiple stages that initiate from floral anlagen; the morphology of FM at this stage is undistinguishable but already shows differential expression of specific marker genes (Smyth et al., 1990; Long and Barton, 2000). The initiation of FM involves many floral integrators including *SUPPRESSOR OF OVEREXPRESSION OF CONSTANS 1* (*SOC1*), which encodes a MADS-box transcription factor that is highly expressed in IM (Lee et al., 2000; Samach et al., 2000). SOC1 directly binds to the promoter of *AGL24* to induce the expression of *AGL24* (Lee et al., 2008). AGL24 also binds to the promoter of *SOC1* to enhance the expression of *SOC1* and form a positive feedback loop for expression (Liu et al., 2008). Meanwhile, the protein complexes formed by the interaction of SOC1 and AGL24 bind to the promoter of *LEAFY* (*LFY*) to enhance *LFY* expression (Lee et al., 2008). The upregulation of *LFY* activates the expression of floral homeotic genes, including *APETALA1* (*AP1*), *CAULIFLOWER* (*CAL*), and *AGAMOUS* (*AG*) to coordinately specify floral organ identities (Blazquez et al., 2006). Upon the induction of *AP1* in the FM, AP1 in turn binds to the promoters of *AGL24*, *SOC1* and another MADS box transcription factor, *SHORT VEGETATIVE PHASE 1* (*SVP1*), to repress the expression of *AGL24* and *SOC1* and prevent the reversion of FM back to IM (Liu et al., 2007). The complex of SOC1 and AGL24 also represses a floral organ identity gene *SEPALLATA3* (*SEP3*) to prevent precocious floral organ development (Gregis et al., 2008; Liu et al., 2009). The suppression of *SOC1* and *AGL24* by AP1 releases the inhibition of *SEP3* to ensure that floral induction and flower development occurs in the proper time and space. In Arabidopsis, floral reversion is observed in transformants ectopically expressing *AGL24*, *SOC1* or *SVP1* (Liu et. al., 2007; 2009), or in the mutants defective in FM identity genes, including *LFY* and *AP1*, or polycomb-group proteins (Okamuro et al., 1996; Müller-Xing et al., 2014). Of note, the expression of *AGL24*, *SOC1* or *SVP1* is detected both in leaf and apex, but whether these floral regulators act as leaf-derived signals in floral reversion remains to be investigated.

Instead of evolving specialized cells to directly sense and transmit perceived signals, plants convert environmental signals into mobile molecules, including mobile mRNAs, and take advantage of vascular systems to distribute mobile mRNAs for inter-organ communication. Plant mobile mRNAs are typical mRNAs that contains a 5’ cap and a polyadenylation tail (Yang and Yu, 2010). To distinguish the mobile from non-mobile mRNAs, the specific motifs or RNA modification on mobile mRNAs can trigger the movement of mobile mRNAs (Zhang et al., 2016; Yang et al., 2019). Mobile mRNAs are selectively targeted to plasmodesmata (PD) for cell-to-cell movement (Luo et al., 2018), or are delivered long distance through phloem to the target tissues (Haywood et al., 2005; Banerjee et al., 2006; Huang et al., 2012; Lu et al., 2012; Huang et al., 2018). Although the detection of translocated mobile mRNAs is corelated with the phenotypic changes in target tissues, whether the translocated mRNAs can be used as templates to synthesize proteins in target tissues remains to be established. However, the proteins encoded by many mobile mRNAs can also move cell-to-cell or long distance (Lucas et al., 1995; Lu et al., 2012), it is technique challenging to distinguish the proteins detected in the target tissues are transported distally or translated locally.

*AGL24* encodes a MADS-box transcription factor that specifies IM differentiation (Yu et al., 2002; 2004). Ectopic expression of *AGL24* promotes floral initiation and IM specification, which leads to a floral reversion phenotype (Yu et al., 2002). The expression of *AGL24* is detected in early developmental stages. In the apex, *AGL24* is detected in IM; however, upon emerging FM, *AGL24* is suppressed by AP1 and is barely detected in FM (Liu et al., 2007). The expression of *AGL24* is not restricted to the shoot apex; it is also detected in vascular tissues of leaves (Yu et al., 2004; Liu et al., 2007; Liu et al., 2008). Interestingly, Arabidopsis grafting experiments showed that *AGL24* mRNA can move long distance from Arabidopsis transformant stocks to the wild-type scions (Yang and Yu, 2010), which suggests that the leaf-expressed *AGL24* mRNA may move to specify the cell fate in FM.

In this study, we provide evidence to show that Arabidopsis *AGL24* mRNA acts non-cell-autonomously to specify meristem identity. Although *AGL24* mRNA was expressed in leaves, AGL24 protein was rapidly degraded and was not accumulated in leaves. The leaf-expressed *AGL24* mRNA can move long distance to the apical meristem where the translocated *AGL24* mRNA can be used as a template to synthesize AGL24 proteins. Sequestering the movement of AGL24 proteins did not abolish the non-cell-autonomous function of *AGL24*, which further supports that protein movement of AGL24 may not explain the non-cell-autonomous function of *AGL24*. Thus, leaf-derived mobile *AGL24* mRNA may provide a systemic signal to bypass transcriptional inhibition by AP1 and convert the cell fate of FM back to IM.

## Results

### Arabidopsis *AGL24* is expressed in leaves during vegetative and reproductive growth

The function of Arabidopsis *AGL24* is to specify IM identity (Yu et al., 2004); however, the expression of *AGL24* is not confined to the apex but rather is mainly detected in leaves (Liu et al., 2007; 2008). Previous Arabidopsis grafting experiments showed that *AGL24* mRNA is mobile when ectopically expressed in leaves (Yang and Yu, 2010), which suggests that the leaf-expressed *AGL24* may act non-cell-autonomously to specify meristem differentiation. Because many mobile genes are expressed in vasculature, which allows the mobile genes to be transported into phloem for spreading throughout the plants (Huang et al., 2012; Chen et al., 2018), we examined the expression pattern of *AGL24* in leaf during vegetative and reproductive stages. A 2.4-kb promoter derived from the promoter region in the *AGL24* genomic DNA fragment (Liu et al., 2007), was PCR-amplified and fused with the β-glucuronidase (*GUS*) reporter gene.

Histochemical analysis of Arabidopsis transformants harboring *AGL24pro*-*GUS* revealed GUS activity mainly in the vasculature of cotyledons, leaves and roots (Figure 1). The detection of *AGL24* promoter activity in vasculature was throughout all developmental stages, from 1-week-old seedlings to mature plants (Figure 1A-D). After floral buds differentiated, the stage when apex-expressed *AGL24* is suppressed by AP1 (Liu et al., 2007), the promoter activity of *AGL24* remained to be detected in leaf vasculature (Figure 1B and 1D). In addition, promoter activity of *AGL24* was detected in root vasculature (Figure 1E). These results indicate that *AGL24* is constitutively expressed in leaf vasculature when apex-expressed *AGL24* is suppressed by AP1.

**Figure 1.**
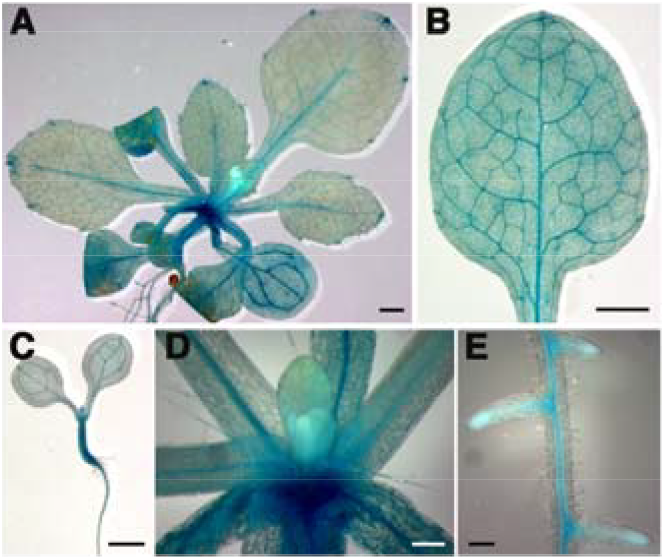
Promoter activity of *AGL24* is detected in vasculature in vegetative and reproductive growth. Histochemical staining of Arabidopsis *AGL24pro-GUS* transformants. (A) 21 day-old transformants, (B) Mature leaves, (C) 7 day-old seedlings, (D) apices and (E) roots. Scale bar (A-C) = 1 mm, (D) = 0.2 mm, (E) = 0.1 mm.

### Leaf-expressed *AGL24* acts non-cell-autonomously to determine cell fate in floral meristem

In Arabidopsis, overexpression of *AGL24* accelerates floral initiation and produces a floral reversion phenotype with a flower emerging in the axials of leaf-like sepals (Yu et al., 2002; 2004). To examine whether leaf-expressed *AGL24* is sufficient to act non-cell-autonomously to regulate floral organ development, the full-length cDNA of *AGL24* was amplified and driven by a control *CaMV35S* promoter or two companion cell (CC)-specific promoters, Arabidopsis *SUCROSE TRANSPORTER 2* (*SUC2*) and pumpkin *GALACTINOL SYNTHASE 1* (*GAS1*). *SUC2* is expressed in CC throughout the plant, whereas *GAS1* is expressed in CC of leaf minor veins (Truernit and Sauer, 1995; Ayre et al., 2003). Arabidopsis transformants harboring *35Spro*-*AGL24*, *SUC2pro*-*AGL24* or *GAS1pro*-*AGL24* showed highly accumulation of *AGL24* mRNA and an early flowering phenotype (Figure 2A and 2B). In addition, the expression of *SEPALLATA3* (*SEP3*), which is downregulated by *AGL24* (Liu et al., 2009), was reduced in the apices of *AGL24* transformants (Figure 2C). These results suggest that CC-expressed *AGL24* can non-cell-autonomously regulate the expression of *SEP3* in apices.

**Figure 2.**
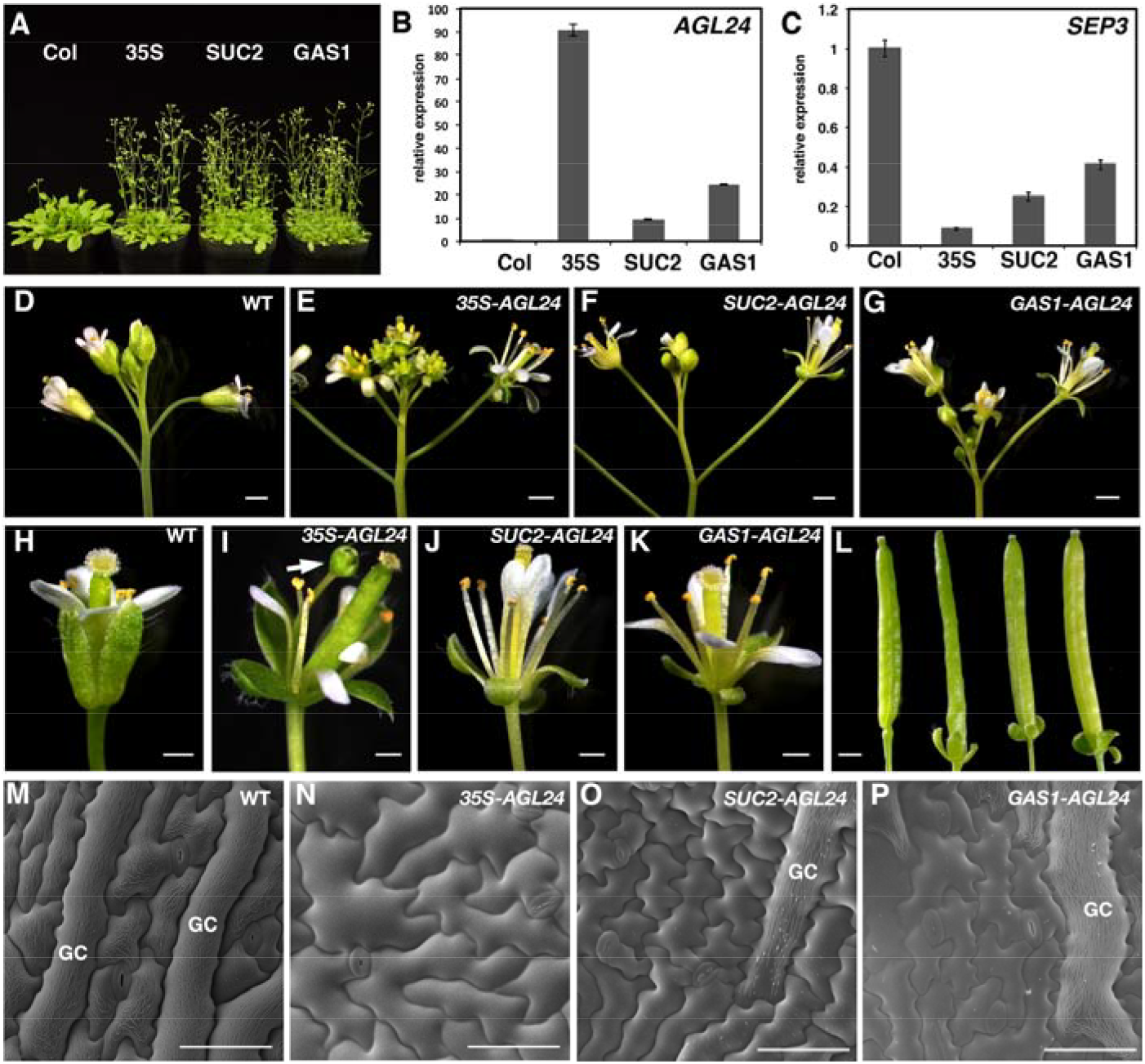
*AGL24* acts non-cell-autonomously to specify floral meristem identity. (A) Flowering time of Arabidopsis wild-type (Col) and *35Spro-AGL24* (35S), *SUC2pro-AGL24* (SUC2), *GAS1-AGL24* (GAS1) transformants. (B, C) Real-time RT-PCR analysis of mRNA levels of *AGL24* (B) or *SEP3* (C) in wild-type (Col) and *35Spro-AGL24* (35S), *SUC2pro-AGL24* (SUC2), *GAS1-AGL24* (GAS1) transformants. The relative expression of *AGL24* or *SEP3* was normalized to β-tubulin expression. (D-G) Floral architecture of Arabidopsis wild-type (D) and *35Spro-AGL24* (E), *SUC2pro-AGL24* (F), *GAS1pro-AGL24* transformants (G). Scale bar= 1000 μm. (H-K) Flower of wild-type (H) and *35Spro-AGL24* (I), *SUC2pro-AGL24* (J), *GAS1pro-AGL24* transformants (K). The extra flower developed from the base of sepals is indicated by an arrow in (I). Scale bar= 500 μm. (L) Siliques of wild-type and *35Spro-AGL24*, *SUCpro2-AGL24*, *GAS1pro-AGL24* transformants (left to right). Note that the leaf-like sepals are attached on the siliques in *AGL24* transformants. Scale bar= 1000 μm. (M-P) Cryo-EM analysis of epidermal cells of sepals in wild-type (M) and *35Spro-AGL24* (N), *SUC2pro-AGL24* (O), and *GAS1pro-AGL24* transformants (P). Note that the giant cells (GCs) are indicated in M, O, P. Scale bar= 50 μm. Data are mean± SD.

The floral organs of *35Spro-AGL24* transformants displayed a typical floral reversion phenotype (Figure 2D-K, Yu et al., 2004). In *35Spro-AGL24* transformants displaying strong floral reversion phenotypes, an extra flower was emerged from the base of sepals (Figure 2I). In addition, the flowers of transformants contained the leaf-like sepals and these sepals remained attached on siliques after flower senescence (Figure 2L). Similarly, *SUC2pro*-*AGL24* or *GAS1pro*-*AGL24* transformants exhibited a comparable phenotype to *35Spro-AGL24* transformants (Figure 2F, 2G); however, the strong floral reversion phenotype was not observed in CC-expressed *AGL24* transformants (Figure 2J, 2K). In addition to floral reversion, the cell identity in floral organs of *AGL24* transformants was changed. On scanning electron microscopy, the epidermal cells of wild-type sepals exhibited a characteristic pattern, with large giant cells interspersed among smaller cells (Figure 2M; Roeder et al., 2010). In contrast, the jigsaw puzzle-shaped cells, which is a characteristic pattern of leaf pavement cells, rather than giant cells were observed in sepals of *35Spro-AGL24* transformants (Figure 2N), which suggests that the FM of *35Spro-AGL24* transformants was converted to IM. In *SUC2pro*-*AGL24* or *GAS1pro*-*AGL24* transformants, sepals contained both giant cells and jigsaw puzzle-shaped pavement cells (Figure 2O-P), which further confirms that CC-expressed *AGL24* acts non-cell-autonomously to convert FM to IM.

### Accumulation of AGL24 proteins was under the detection limit in Arabidopsis leaves

To investigate whether the non-cell-autonomous function of *AGL24* is attributed to the movement of AGL24 proteins, we examined the accumulation of AGL24 proteins in leaves. We tagged AGL24 with GREEN FLUORESCENT PROTEIN (GFP) and expressed *GFP-AGL24* fusion constructs in Arabidopsis. Transformants harboring *35Spro-GFP-AGL24* or *SUC2pro-GFP-AGL24* displayed an *AGL24* overexpression phenotype (Supplemental Figure 1), suggesting that the fusion with GFP did not affect the function of AGL24. Unlike *35Spro-EGFP* or *SUC2pro-EGFP* transformants, with GFP fluorescence detected in leaves (Figure 3A, 3C), GFP fluorescence was absent in leaves of transformants harboring *35Spro-GFP-AGL24* or *SUC2pro-GFP-AGL24* (Figure 3B, 3D). On western blot analysis with antibodies against Arabidopsis AGL24, endogenous AGL24 or GFP-AGL24 fusion protein was under the detection limit in leaf (Figure 3E), but AGL24 or GFP-AGL24 fusion protein was detected in apices of transformants (Figure 3E), which indicates that AGL24 proteins were not accumulated in leaves where *AGL24* mRNA was expressed. Thus, the non-cell-autonomous function of *AGL24* in *SUC2pro-GFP-AGL24* transformants is not likely explained by the movement of AGL24 protein from leaves. This notion is further supported by the expression of movement-defective DsRED-AGL24 fusion protein in Arabidopsis. DsRED is a tetrameric RFP version that has been used to limit the movement of mobile proteins (Lu et al., 2012). Arabidopsis transformants carrying *SUC2pro-DsRED-AGL24* still displayed a phenotype resembling *AGL24*-overexpressed lines (Supplemental Figure 2). In agreement with the absence of AGL24 protein in leaves, the expression of *SOC1*, which is directly activated by AGL24 (Liu et al., 2008), was induced in apices but not leaves of *35Spro-GFP-AGL24* or *SUC2pro-GFP-AGL24* transformants (Figure 3F and 3G), suggesting that *AGL24* did not activate downstream gene expression in leaves. Consistent with this notion, we observed no obvious vegetative phenotypes in *agl24* or *agl24*, *soc1*, *svp* triple mutants (Supplemental Figure 3), although previous study observed a phenotype of late flowering in *agl24* or premature transition of IM to FM in which the chimeric floral structure and an extra bract subtends each floral structure in *agl24*, *soc1*, *svp* triple mutants (Liu et al., 2009). Taken together, our results indicate that the non-cell-autonomous function of *AGL24* may not be attributed to the movement of AGL24 protein or the downstream products activated by *AGL24* in leaves.

**Figure 3.**
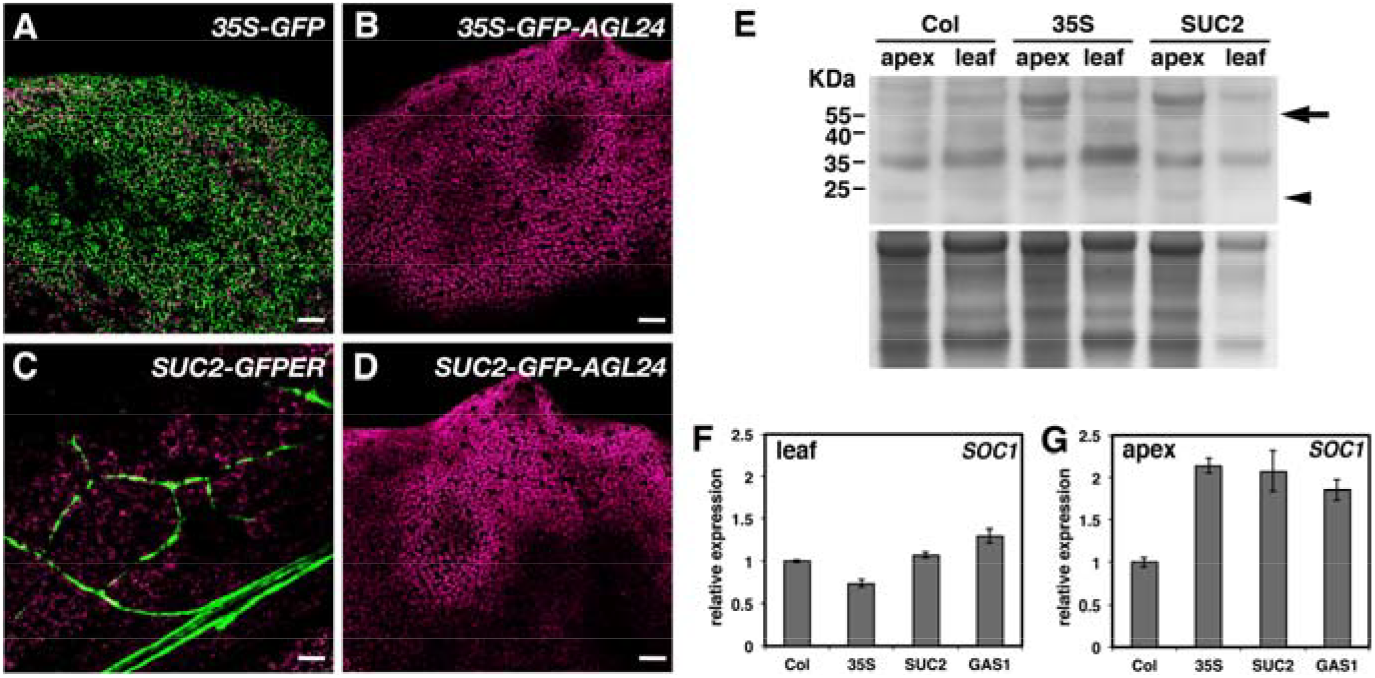
Accumulation of AGL24 proteins was under the detection limit in Arabidopsis leaves. (A-D) Confocal microscopy of GFP fluorescence in leaves of Arabidopsis *35Spro-GFP* (A), *SUC2pro-GFP-ER* (B), *35Spro-GFP-AGL24* (C), and *SUC2pro-GFP-AGL24* (D) transformants. Scale bar= 50 μm. (E) Immunoblot assay of proteins extracted from apex or leaf of wild type (Col) and *35Spro-GFP-AGL24* (35S) or *SUC2pro-GFP-AGL24* (SUC2) transformants. Upper panel, antibodies against AGL24 were used for immunoblot assay. Proteins of GFP-AGL24 or AGL24 are indicated by an arrow and arrowhead, respectively. The protein size marker is indicated (kDa). Lower panel, the Coomassie blue-stained loading control. (F, G) Real-time RT-PCR analysis of *SOC1* mRNA level in leaves (F) or apex (G) of wild-type (Col) and *35Spro-AGL24* (35S), *SUC2pro-AGL24* (SUC2), and *GAS1-AGL24* (GAS1) transformants. The expression of *SOC1* was normalized to β-tubulin. Data are mean± SD.

### AGL24 protein encounters rapid degradation in leaves

To verify whether no detection of AGL24 protein in leaves is owing to rapid degradation of AGL24 protein in leaves, we treated *35Spro-GFP-AGL24* or *SUC2pro-GFP-AGL24* transformants with MG132, a 26S proteasome inhibitor (Lee and Goldberg, 1998). Confocal microscopy detected GFP fluorescence in the nucleus of *35Spro-GFP-AGL24* transformant leaves, indicating that proteosome inhibitor treatment reduced AGL24 protein degradation in leaves (Figure 4A, 4B). AGL24 protein belongs to MIKC-type MADS-box transcription factors that contains an N-terminal MADS (M) domain, an intervening (I) domain, a keratin-like (K) domain, and a C-terminal (C) domain (Theiβen et al., 1996). To examine the domain involved in AGL24 protein degradation in leaves, we PCR-amplified *AGL24* fragments containing different domains and fused these fragments with GFP for transient assay (Figure 4C-H). In control experiments, Agroinfiltration revealed strong GFP fluorescence in *Nicotiana benthamiana* leaves expressing *35S-GFP* but no GFP fluorescence in leaves expressing *35S-GFP-AGL24* (Figure 4C, 4D). When the M-domain was truncated from AGL24 (GFP-IKC), GFP fluorescence was restored (Figure 4F), which suggests that the M-domains is required for AGL24 protein degradation in leaves. In addition, the M-domain is sufficient to mediate AGL24 protein degradation in leaves because the fusion of GFP with the M-domain (GFP-M), but not the C-domain of AGL24 (GFP-C) resulted in no detection of GFP fluorescence in leaves (Figure 4G and H). These results indicated that the M-domain of AGL24 is necessary and sufficient to trigger AGL24 protein degradation in leaves.

**Figure 4.**
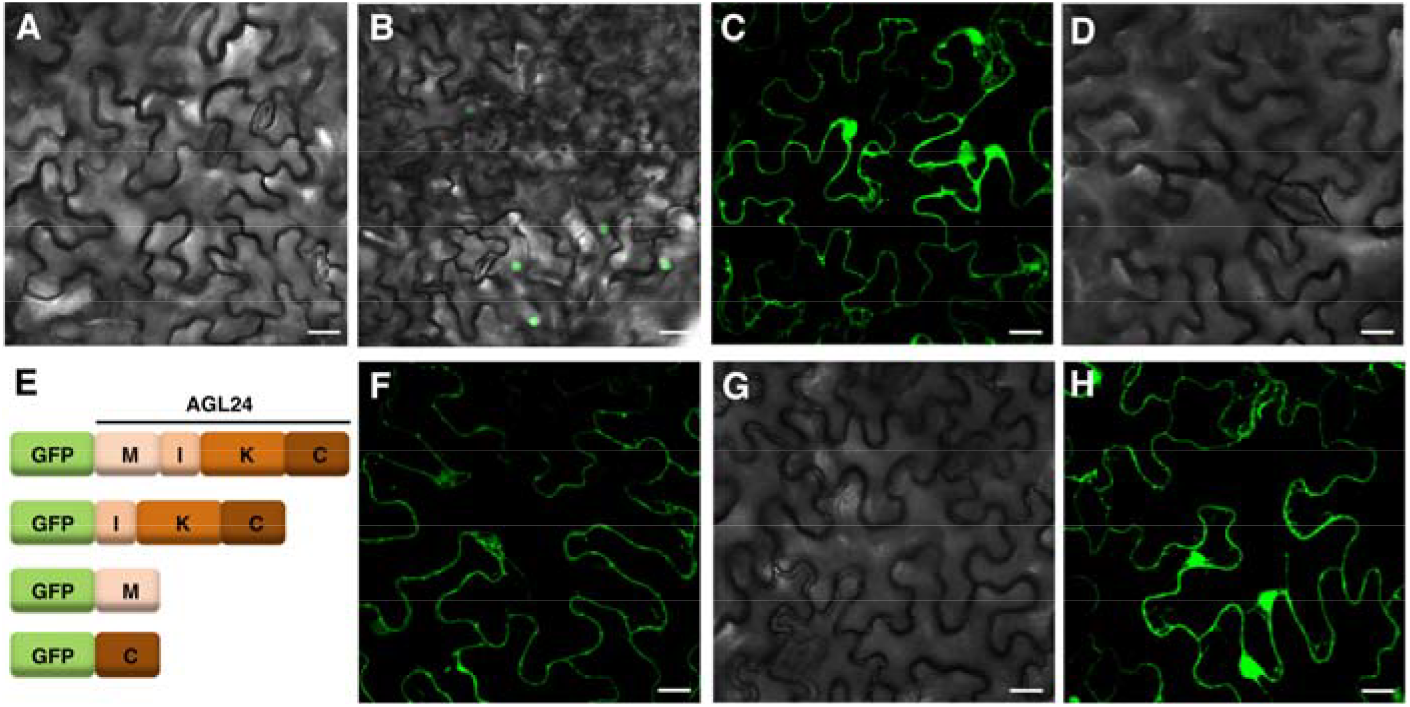
AGL24 proteins encounter rapid degradation in leaves. (A, B) Confocal microscopy of Arabidopsis *35Spro-GFP-AGL24* transformants treated with DMSO (A), or MG132 (B) for 16 h. Scale bar= 20 μm. (C-H) Truncation analysis identified motifs required for AGL24 protein degradation in leaves. Transient expression of *35Spro-GFP* (C), *35Spro-GFP-AGL24* (D), in *N. benthamiana* leaves. (E) Illustration of *35Spro-GFP-AGL24* motifs. Transient expression of *35Spro-GFP-IKC* (F), *35Spro-GFP-M* (G), or *35Spro-GFP-C* (H) motifs in *N. benthamiana* leaves. Scale bar= 20 μm.

### Translocated *AGL24* mRNAs can serve as a template for protein translation

We have previously shown that *AGL24* mRNA can move long-distance from ectopically expressed transformant stocks to wild-type scions (Yang and Yu, 2010). To examine whether endogenous *AGL24* mRNA is a mobile mRNA, we grafted Arabidopsis *agl24* null mutants onto wild-type stocks. We obtained a T-DNA insertion mutant, *agl24-3* (Salk_095007C), with a T-DNA inserted at exon 4 of *AGL24*. The level of *AGL24* mRNA in *agl24-3* was under the detection limit, indicating that *agl24-3* is a null mutant (Figure 5A). At 2 weeks after grafting, real-time RT-PCR detected *AGL24* mRNA in *agl24-3* scions grafted onto wild-type stocks but not from *agl24-3* scions grafted onto *agl24-3* stocks (Figure 5B), which indicates that endogenous *AGL24* mRNA moved long distance across the graft union. To further examine whether *AGL24* mRNA can trigger the movement of a non-mobile mRNA, we used Arabidopsis transformants expressing *GFP-AGL24* chimeric mRNA for grafting. At 2 weeks after grafting, *GFP-AGL24* mRNA was detected in the wild-type scions grafted onto *35Spro-GFP-AGL24* or *SUC2pro-GFP-AGL24* transformant stocks. (Figure 5C-D), which supports that *AGL24* mRNA can trigger the movement of a non-mobile mRNA.

**Figure 5.**
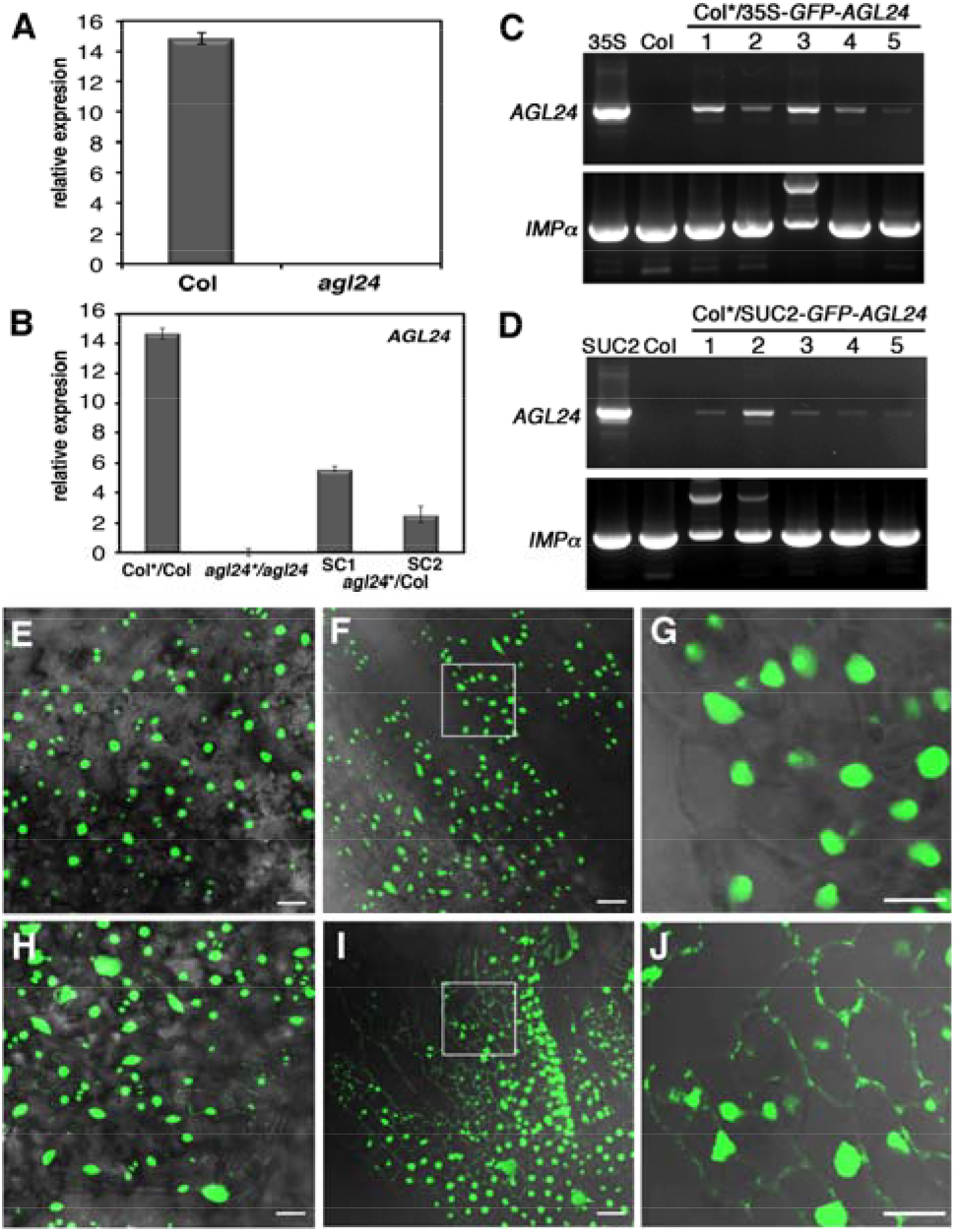
*AGL24* mRNA is a mobile mRNA. (A, B) Endogenous *AGL24* mRNA movement in Arabidopsis grafting experiments. Real-time RT-PCR analysis of *AGL24* mRNA in wild-type (Col) and *agl24-3* (*agl24*) null mutants (A); or scions (*) of Arabidopsis grafting experiments (B). Total RNA was extracted from wild-type scions grafted onto wild-type stocks (Col*/Col); *agl24-3* scions onto *agl24-3* stocks (*agl24*\*/*agl24*), *agl24-3* scions onto wild-type stocks (*agl24*\*/Col). SC1 and SC2 are two independent scions in *agl24-3* scions grafted onto wild-type stocks. Relative expression of *AGL24* was normalized to *UBIQUITIN CONJUGATING ENZYME*. (C-D) Chimeric *GFP-AGL24* mRNA is mobile in Arabidopsis grafting experiments. RT-PCR analysis of mRNA level in wild-type scions (Col*) grafted onto (C) *35Spro-GFP-AGL24* transformant stocks or (D) *SUC2pro-GFP-AGL24* transformant stocks. RNA of wild-type (Col), *35Spro-GFP-AGL24* (35S), or *SUC2pro-GFP-AGL24* (SUC2) was used as controls. Five independent scions were used for RT-PCR. *IMPORTIN-α* (*IMP-α*) was used as a loading control. Data are mean± SD. (E-J) MS2-based mRNA live-imaging of Arabidopsis transformants carrying *35Spro-FD_NLS_-MS2-GFP* (E-G) or the scion tissues from *35Spro-FD_NLS_-MS2-GFP* plants that grafted onto *35Spro-AGL24_SL48_* stocks (H-J). (E, H) Cauline leaf; (F, I) Sepal; (G, J) The magnified image of the highlight area from F, I.

To examine whether *AGL24* mRNA is transported to the apical meristem to specify floral organ differentiation, we employed an MS2-based mRNA live-imaging system to directly visualize translocated *AGL24* mRNA in scions. This system has been used to visualize selective PD-targeting of mobile *AGL24* mRNA (Luo et al., 2018). The MS2 system employed a nuclear-localized and GFP-fused coat protein of bacteriophage MS2 (FD_NLS_-MS2-GFP). When the FD_NLS_-MS2-GFP fusion protein was co-expressed with target mRNA conjugated to multiple copies of the stem-loop structure (SL) that specifically recognizes by MS2 coat protein, the localization of target mRNA was revealed by binding of FD_NLS_-MS2-GFP to SL-containing target mRNAs (Luo et al., 2018). We generated Arabidopsis transformants carrying *35Spro-FD_NLS_-MS2-GFP* or *35Spro-AGL24_SL48_*. On confocal microscopy of *35Spro-FD_NLS_-MS2-GFP* transformants, GFP fluorescence was restricted in the nucleus (Figure 5E-G). After grafting *35Spro-FD_NLS_-MS2-GFP* scions onto *35Spro-AGL24_SL48_* stocks, confocal microscopy detected GFP fluorescence in the cell periphery of sepal, petal and cauline leaves (Figure 5H-J). The GFP fluorescence detected in the cell periphery displayed punctate-like foci (Figure 5J), which is consistent with previous live imaging analysis of *AGL24* mRNA in transient assay (Luo et al., 2018).

To investigate whether translocated *AGL24* mRNA can serve as a template for protein translation in apices, we examined the floral apices of *SUC2pro-GFP-AGL24* transformants and detected GFP fluorescence in the nucleus of floral apices (Figure 6A and 6B). To verify that GFP fluorescence in apex was locally translated from translocated *GFP-AGL24* mRNA, we applied the translational inhibitor cycloheximide onto the apices of *SUC2pro-GFP-AGL24* transformants. At 2 h after cycloheximide treatment, GFP fluorescence was greatly reduced in apices of *SUC2pro-GFP-AGL24* transformants (Figure 6C), so the translocated *GFP-AGL24* mRNA can be used as a template to translate into proteins.

**Figure 6.**
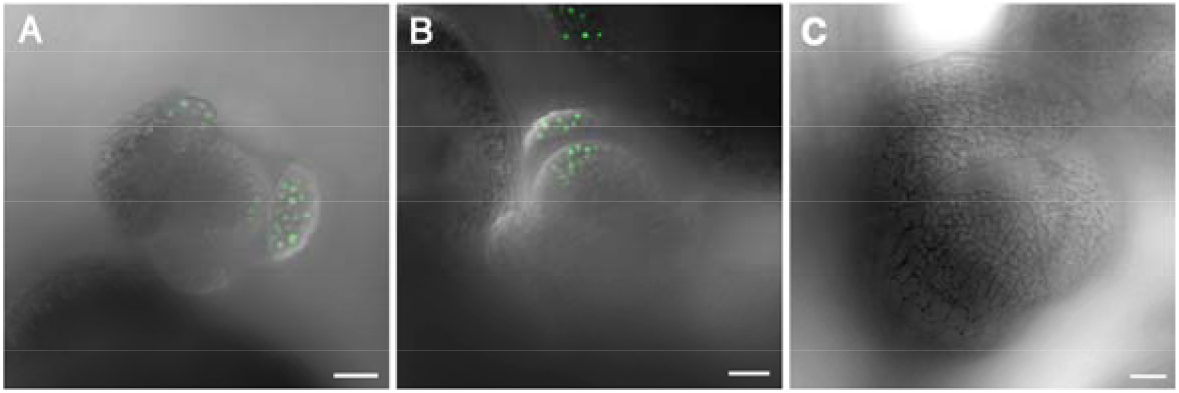
Translocated *GFP-AGL24* mRNA is used as a template for protein translation in apex. GFP fluorescence was detected in floral apices of *35Spro-GFP-AGL24* (A) or *SUC2pro-GFP-AGL24* transformants (B). (C) Cycloheximide treatment abolished the detection of GFP fluorescence in floral apices of *SUC2pro-GFP-AGL24* transformants. Scale bar= 20 μm.

### Leaf-expressed *AGL24* is involved in floral organ development

To further verify whether vasculature-expressed *AGL24* is required for FM specification, we specifically knocked down *AGL24* expression by artificial microRNA against *AGL24* (*AGL24-amiR*). In contrast to systemic RNAi, amiRNA has a limited non-cell-autonomous effect (Schwab et al., 2006; McHale et al., 2013). In addition, the long-distance movement of amiRNA from vasculature to the apical meristem is restricted (Skopelitis et al., 2018), which allows for knocking down the expression of *AGL24* in specific tissue. Given that the triple mutant *agl24*, *soc1*, *svp* shows a floral organ defect phenotype (Liu et al., 2009), we introduced *SUC2-* or *GAS1*-*AGL24-amiR* to the *soc1, svp* double mutant. In control experiments, Arabidopsis *soc1, svp* transformants harboring *35Spro*-*AGL24-amiR* or *35Spro*-*AGL24-IR* (the inverted repeat of *AGL24* that induced systemic RNAi) showed a phenotype with premature transition of IM to FM that was reminiscent of the *agl24*, *soc1*, *svp* triple mutants (Liu et al., 2009; Figure 7A-D).

**Figure 7.**
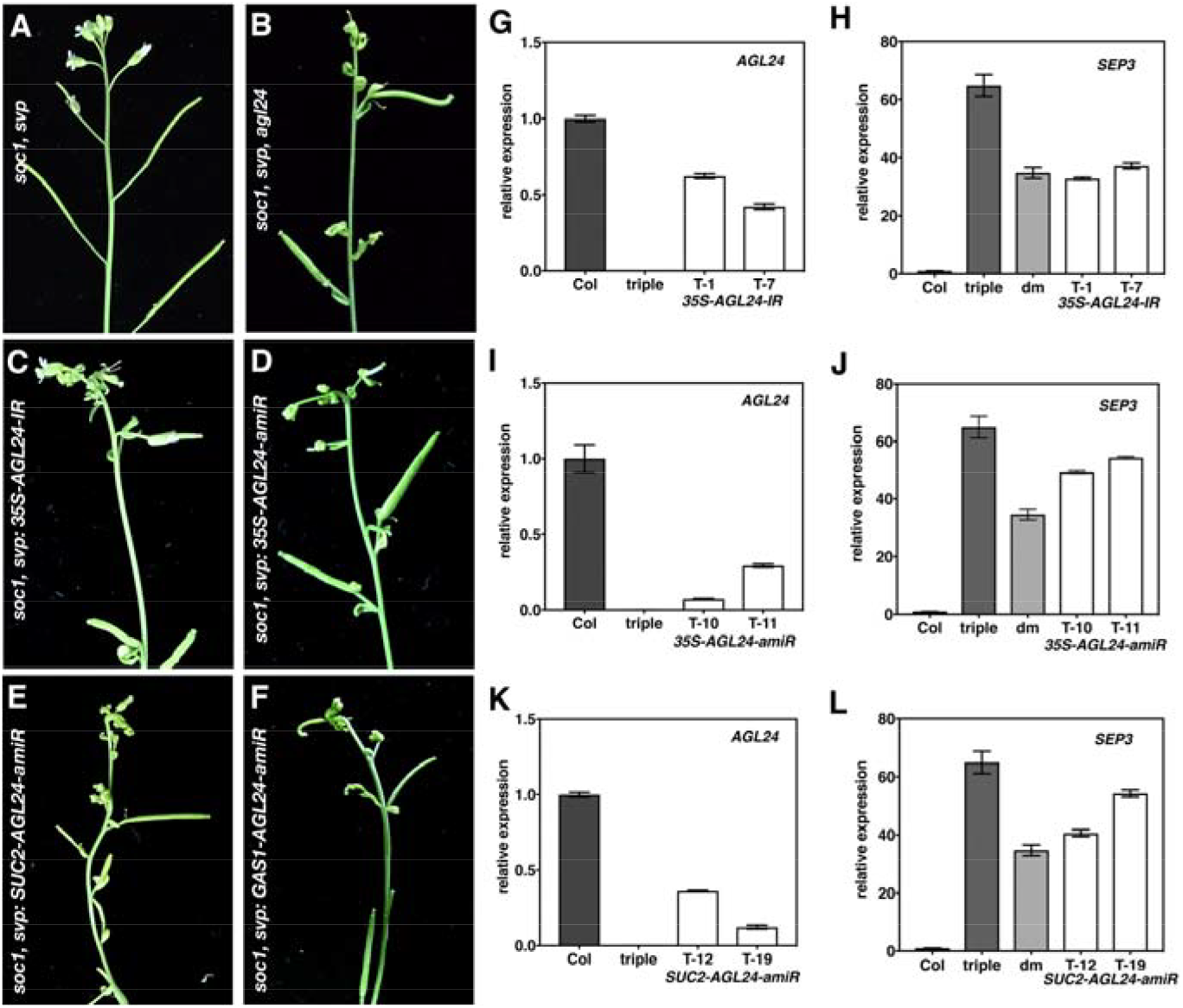
Expression of *AGL24* in leaf is required for floral meristem development. (A-F) Floral architecture of *soc1*, *svp* double mutant (A), *soc1*, *svp*, *agl24-3* triple mutant (B), or *soc1*, *svp* double mutant harboring *35Spro-AGL24-IR* (C), *35Spro-AGL24-amiR* (D), *SUC2pro-AGL24-amiR* (E), or *GAS1pro-AGL24-amiR*. (G-L) Real-time RT-PCR analysis of *AGL24* or *SEP3* mRNA in *soc1*, *svp* double mutant harboring *35Spro-AGL24-IR* (G, H), (I, J) *35Spro-AGL24-amiR*, or (K, L) *SUC2pro-AGL24-amiR*. Total RNA extracted from wild-type (Col), *soc1*, *svp*, *agl24-3* triple mutant (triple), *soc1*, *svp* double mutant (dm), two representative transformants was used for analysis. Relative expression of *AGL24* and *SEP* was normalized to *UBIQUITIN CONJUGATING ENZYME*. Data are mean± SD.

Similarly, transformants harboring CC-expressed *AGL24-amiR* showed a triple mutant phenotype (Figure 7E and F). Consistent with phenotypic alterations, the accumulation of *AGL24* mRNA in various *AGL24-IR* or *AGL24-amiR* transformants was reduced (Figure 7G, 7I, 7K). In addition, the expression of *SEP3* was upregulated in apices of these transformants (Figure 7H, 7J, 7L). These results suggest that knocking down the expression of *AGL24* in companion cells can non-cell-autonomously affect FM differentiation.

## Discussion

The use of mRNA as a mobile molecule is an efficient approach to distribute proteins to specific locations. Instead of translocating quantity proteins, the transport of mRNA coupled with local translation is relatively energy-conserving. In addition, upon translocating to target cells, mRNAs may continuously serve as templates to synthesize proteins even though the environmental conditions that induce mobile mRNA expression no longer exist. Thus, the translocated mobile mRNAs may prolong gene induction by temporarily storing the environmental inputs in genetic codes to strengthen the perceived signals. Although the use of heterospecies grafting has identified many plant-mobile mRNAs (Thieme et al., 2015; Yang et al., 2015; Notaguchi et al., 2015; Huang et al., 2018; Xia et al., 2018), whether these mobile mRNAs indeed act as systemic signals remains controversial. Unlike many mobile mRNAs in which the proteins encoded by mobile mRNAs are also mobile, the proteins of AGL24 are rapidly degraded in leaves (Figure 3, 4), which may limit the mobility of AGL24 proteins from leaves. Restriction of protein movement by fusing AGL24 with DsRED did not affect the non-cell-autonomous function of *AGL24* (Supplemental Figure 2), which further supports that the non-cell-autonomous function of *AGL24* is not attributed to the movement of AGL24 proteins. In addition, ectopically expressed *AGL24* in leaves activated *SOC1* in the apex but not leaves (Figure 3F, 3G), which is consistent with the absence of AGL24 proteins in leaves. Thus, the non-cell-autonomous function of *AGL24* is not likely attributed to the movement of AGL24 proteins or the downstream gene products of *AGL24* but rather is caused by the movement of *AGL24* mRNA. Indeed, *AGL24* is expressed in vasculature (Figure 1), which is consistent with the detection of *AGL24* mRNA in broccoli phloem sap (Yang and Yu 2010). Grafting experiments demonstrated that *AGL24* mRNA is a mobile mRNA (Figure 5). In addition, the translocated *AGL24* mRNA can be used as a template for protein translation in the apex (Figure 6). Thus, mobile *AGL24* mRNA can move from leaves to apices and serve as a template for protein translation to specify meristem development.

Floral organ development is one of the critical developmental steps for plants to produce progenitors. To maximize seed production, plants sense and convert the environmental dynamics into mobile molecules to synchronize floral development with seasonal clues. Indeed, to fine-tune the transition of VM to IM, photoperiodic variations trigger a pair of antagonized mobile molecules, florigen and antiflorigen, which are encoded by homologues with opposite function (Corbesier et al., 2007; Lu et al., 2012; Huang et al., 2012). Similarly, the movement of *AGL24* from leaf to apex may reflect environmental variations. It has been shown that the expression of *AGL24* is affected by light, hormone and vernalization (Yu et al., 2002; Michaels et al., 2003). During FM establishment, AP1 binds to the promoter of *AGL24* to prevent continued of IM development (Liu et al., 2007). The movement of *AGL24* mRNA from leaves may provide another layer of post-transcriptional control to bypass AP1 transcriptional inhibition and adjust FM back to IM development. In Arabidopsis, *AGL24* and SVP are in the same clade in the MADS gene family (Martínez-Castilla et al. 2003). Whether SVP or other MADS genes also act as a systemic signal remains to be investigated.

The treatment of Arabidopsis 35S-GFP-AGL24 transformants with proteosome inhibitor MG132 suggests that AGL24 proteins are rapidly degraded in leaves (Figure 4). MG132 inhibits the proteolytic activity of 26S proteasomes, which is required for the degradation of ubiquitin conjugation proteins (Lee and Goldberg, 1998). This observation suggests that ubiquitination may be involved in the rapid degradation of AGL24 proteins in leaves. The prediction of ubiquitination of AGL24 proteins demonstrates that the putative ubiquitination sites are located at the C-terminal sequences; however, our analysis showed that the truncation of the C-terminal from AGL24 proteins did not affect protein stability. Instead, the M-domain is necessary and sufficient to mediate AGL24 protein degradation (Figure 4), suggesting that ubiquitination may indirectly regulate AGL24 protein stability. The M-domain in AGL24 protein is required for DNA binding and is involved in dimerization and nuclear localization (Theiβen et al., 2016). It is possible that the protein-protein interaction of AGL24 and different MADS-box transcription factors may stabilize AGL24 proteins. Consistent with this hypothesis, recent results indicate that differential dimerization of MADS-box transcription factors in maize may affect protein degradation (Abraham-Juárez et al., 2020). In addition to post-translational control of AGL24 proteins in leaves, our results did not preclude that the leaf-expressed *AGL24* mRNA may also encounter translational control to limit protein translation. Intra-and inter-cellular targeting of mobile mRNA to PD for cell-to-cell movement is a highly regulated process. However, the translational complex assembled by ribosomes and mRNA is a huge protein complex that oversizes the size exclusion limit of PD. How mobile mRNAs escape from sequestration by ribosome complexes and successfully transport through PD remains to be elucidated.

The use of amiRNAs is an effective approach to specifically knock down the expression of target genes in a tissue-specific manner. Although grafting experiments have shown that plant small RNAs, including small interfering RNAs (siRNAs) and miRNAs, have the ability to move cell-to-cell and long-distance (Liu and Chen 2018; Maizel et al., 2020), the modes of siRNAs and miRNAs movement differ. In contrast to siRNAs, which spread and trigger systemic silencing in a wide range, the movement of miRNAs and amiRNAs is a highly regulated process and has relatively limited non-cell-autonomous effects (Schwab et al., 2006; de Feloppes et al., 2011; Skopelitis et al., 2018). Our results with CC-expressed *AGL24-amiR* in the *soc1, svp* double mutant revealed a triple mutant phenotype, suggesting that knocking down the expression of *AGL24* in leaves may affect FM differentiation (Figure 7). This result is consistent with the notion that mobile *AGL24* mRNA is a leaf-derived signal to regulate floral organ development. In *Impatiens balsamina*, defoliation experiments showed that floral reversion is controlled by a leaf-derived signal (Pouteau et al., 1997; Tooke and Battey, 2000). Analysis of *Impatiens* homologs of *AP1* and *LFY* suggested that these two genes may not be the leaf-derived signals to control floral reversion (Pouteau et al., 1997; Tooke et al., 2005). Whether the mRNA movement of an *Impatiens* homolog of *AGL24* can control floral reversion remains to be investigated.

## Methods

### Plant materials and growth conditions

*Arabidopsis thaliana agl24-3* (salk_095007C) and *svp-41* seeds were obtained from Arabidopsis Biological Resource Center (ABRC, Ohio, USA). The double mutant of *soc1*, *agl24* was provided by Dr. Hao Yu (Department of Biological Sciences, National University of Singapore). The triple mutant of *soc1*, *agl24*, *svp-41* was generated by crossing *soc1*, *agl24* with *svp-41*. The plants were grown in growth chambers under 16/8-h, 22^°^C/20^°^C day/night cycle, under white fluorescent light, at light intensity of 100 μmol m^−2^ s^−1^.

### Plasmid construction

The sequences of the gene-specific primers used to amplify the promoter of *AGL24* or full-length *AGL24* cDNA are in Supplemental Table S1. To generate *AGL24* truncation or AGL24-IR clones, the fragments of *AGL24* were PCR-amplified with gene-specific primers in Supplemental Table S1. The artificial miRNA (amiRNA) was generated according to Schwab et. al., 2006. The primers used to generate *AGL24* amiRNA were designed based on WMD3 prediction (http://wmd3.weigelworld.org). The sequences of the primers are in Supplemental Table S1. All constructs were confirmed by sequencing. Arabidopsis transformants were selected on MS medium containing 40 μg/mL hygromycin. The antibiotic-resistant plants were transferred to soil for further analysis.

### GUS staining

For GUS activity assay, Arabidopsis transformants were incubated with GUS staining solution (50 mM sodium phosphate pH 7.0, 10 mM EDTA, 0.5 mM potassium ferricyanide, 0.5 mM potassium ferrocyanide, 1 mM X-Gluc, 0.01% Triton X-100) at 37^°^C for 16 h. Plants were treated with 95% ethanol to remove chlorophyll and photographed under a dissect microscope.

### Arabidopsis seedling grafting

Arabidopsis seedling grafting experiments followed previous procedures (Huang and Yu, 2015). In brief, 10-to 12-day-old soil-grown plants were used for grafting. The apexes of stocks were removed by using a microdissecting scissor under a dissecting microscope. The scions were cut from hypocotyls, and the mature leaves of scions were removed from 1/3 of the leaf blade. The hypocotyls of the scions were cut from 0.1 cm below the cotyledons by using a double-edged stainless steel razor blade to create a flat surface. To assemble the scions with rootstocks, a 0.1-mm insect pin was inserted into scions from the base of the petiole through the hypocotyl. The pin-containing scions were vertically attached onto the apex-removed rootstocks. The grafted plants were kept in a covered tray to retain humidity for 1 week, then the lid was removed and plants were transferred to normal growth conditions. Two weeks after grafting, the scions were subjected to RT-PCR or confocal microscopy analyses.

### RNA extraction and RT-PCR analysis

Total RNA from plant tissues was extracted by use of TRIzol reagent (Invitrogen). For RT-PCR analysis, 5 μg total RNA was used in reverse-transcription reactions performed with oligo(dT)_20_ and SuperScript III reverse transcriptase (Invitrogen). A 1 μl amount of cDNA was used for the PCR reaction with the following conditions: 1 min at 94^°^C for 1 cycle; 30 sec at 94^°^C, 30 sec at 60^°^C, 1min at 68^°^C for 35 cycles and 10 min at 68^°^C for 1 cycle. Specific primers against the transgene and *NOS* terminator were used for PCR reactions to distinguish transgenes from endogenous genes (primer sequences are in Table S1). To minimize the variations in PCR, the same primers were used for each set of experiments. PCR was conducted at least twice for each sample to make sure the data is representative. An aliquot (5 μl) of PCR products was separated on 1.5% agarose gels.

### Protein extraction and immunoblotting analysis

Arabidopsis total proteins were extracted from the floral apex or mature leaves of wild-type or *AGL24* transformants with protein extraction buffer (0.3 mM Tris-HCl pH8.5, 8% SDS, 1 mM EDTA, and 1X complete protease inhibitor cocktail). The samples were boiled in 100°C water for 5 min before separating on SDS-PAGE.

For immunoblot analysis, 10 μg of total proteins was separated on SDS-PAGE (12% NuPAGE Novex Bis-Tris mini gels, Invitrogen) with 1X NuPAGE 2-(N-morpholino) ethanesulfonic acid (MES) buffer. The separated proteins were transferred to polyvinylidene difluoride (PVDF) membrane. Membranes were incubated in blocking reagent [5% non-fat milk in 1X TBS (20 μM Tris-HCl pH 7.4, 150 mM NaCl)] at room temperature for 30 min. After 3 times washed with 1X TBS, membranes wwere incubated with 1:1000 diluted rabbit anti-AGL24 antibodies. The ABC Peroxidase Staining Kit (Thermo Scientific) with the substrate solution (6 mg diaminobenzidine tetrachloride, 8 mg NiCl_2_, and 10 μl H_2_O_2_ in 10 ml 1XTBS) was used to detect the protein signals. The SDS-PAGE stained with Coomassie blue was used as a loading control.

### Cryo-scanning electron microscopy

Floral tissues were cut and loaded on a cryo-specimen holder and frozen in liquid nitrogen. The holder with frozen tissues was transferred to a sample preparation chamber at −160 ^°^C for 5 min, with temperature increased to −85 ^°^C for 15 min. After being coated with platinum (Pt) at −130 ^°^C, samples were transferred to a cryo stage in an SEM chamber and observed at −160 ^°^C by use of a cryo-scanning electron microscope (FEI Quanta 200 SEM/Quorum Cryo System PP2000TR FEI) at 20KV.

### Confocal laser scanning microscopy

GFP fluorescence was observed under a confocal laser scanning microscopy LSM880 (Carl Zeiss) with the excitation/emission filter sets of argon 488/500-530 nm.

### MG132 or cycloheximide treatment

For MG132 treatment, 3-week-old *35Spro-GFP-AGL24* plants were cut from the hypocotyls and soaked in Murashige and Skoog medium containing 50 μM MG132 or DMSO for controls. The plants were incubated at room temperature with shaking for 16 h. GFP fluorescence was examined under confocal laser scanning microscopy.

For cycloheximide treatment, the floral apices of 5-week-old Arabidopsis *SUC2pro-GFP-AGL24* transformants were dipped into cycloheximide solution (50 μM cycloheximide, 0.015% Silwet L-77, 0.1% EtOH) for 3-5 sec. At 2 h after dipping, GFP fluorescence was examined under a confocal laser scanning microscopy.

## Supplemental Data

**Supplemental Figure 1**. Arabidopsis *GFP-AGL24* transformants displayed an *AGL24* overexpression phenotype.

**Supplemental Figure 2**. Arabidopsis *SUC2pro-DsRED-AGL24* transformants displayed an *AGL24* overexpression phenotype.

**Supplemental Figure 3**. Arabidopsis *agl24* or *agl24*, *soc1*, *svp* triple mutants displayed no obvious vegetative phenotypes.

**Supplemental Table 1**. Primers used in RT-PCR and real-time RT-PCR analysis

## Acknowledgements

We thank Dr. Hao Yu (Department of Biological Sciences, National University of Singapore) and the Arabidopsis Biological Resource Stock Center (ABRC) for providing Arabidopsis seeds and Dr. Wan-Nan Jane for providing technique support on cryo-SEM. We also thank Yu-Shan Liu for the characterization of *AGL24* transformants and mutants. This work was supported by grants from the Ministry of Science and Technology,

## Author Contributions

The experiments were designed and done by N.C.H. and T.S.Y. GUS staining and Western blot were done by H.C.T. The manuscript was written by T.S.Y. with the help of N.C.H. and H.C.T.

